# Epithelial galectin-3 induces mitochondrial complex inhibition and cell cycle arrest of CD8^+^ T Cells in severe/critical ill COVID-19

**DOI:** 10.1101/2023.03.14.532609

**Authors:** Yudie Wang, Cheng Yang, Zhongyi Wang, Yi Wang, Qing Yan, Ying Feng, Yanping Liu, Xiaolan Zhang, Jingwei Zhao, Juan Huang, Jingjiao Zhou

**Affiliations:** Department of Biology and Genetics, College of Life Sciences and Health, Wuhan University of Science and Technology, Wuhan, China; Department of Medicine, Maternal and Child Health Hospital, Tongji Medical College, Huazhong University of Science and Technology, Wuhan, China

**Keywords:** COVID-19, CD8^+^ T cell, Mitochondrial complex genes, Cell cycle arrest, Galectin-3

## Abstract

Several studies have identified the presence of functionally depleted CD8^+^ T cells in COVID-19 patients, and particularly abnormally reduced CD8^+^ T cells in severe/critical patients, which may be a major cause of disease progression and poor prognosis. In this study, a proliferating-depleted CD8^+^ T cell phenotype was observed in severe/critical COVID-19 patients through scRNA-seq and scTCR-seq analysis. These CD8^+^ T cells were subsequently found to be characterized by cell cycle arrest and downregulation of mitochondrial biogenesis and respiratory chain complex genes. Cellchat analysis revealed that the Galectin signaling pathways between infected lung epithelial cells and CD8^+^ T cells play the key role in inducing CD8^+^ T cell reduction and dysfunction in severe/critical COVID-19. We used SARS-COV-2 ORF3a to transfect A549 epithelial cells, and co-cultured with CD8^+^ T cells. The ex vivo experiments confirmed that galectin-3 inhibited the transcription of mitochondrial respiratory chain complex III/IV genes in CD8^+^ T cells by suppressing the nuclear translocation of nuclear respiratory factor 1 (NRF1). In addition, the regulatory effect of galectin-3 was correlated with the activation of ERK signaling and/or the inhibition of Akt signaling. Galectin-3 inhibitor, TD-139, promoted nuclear translocation of NRF1, and enhanced mitochondrial respiratory chain complex III/IV gene expression and mitochondrial biogenesis, then restore the expansion ability of CD8^+^ T cells. Our study improved the understanding the immunopathogenesis and provided new target for the prevention and treatment of severe/critical COVID-19.

## Introduction

Severe acute respiratory syndrome coronavirus 2 (SARS-CoV-2) emerged in late 2019 caused pandemic of acute respiratory disease, named ‘coronavirus disease 2019’ (COVID-19). Emerging variants of SARS-CoV-2 are continuing to ravage many countries, leading to reinfections and breakthrough infections, causing repeated outbreaks, which have taken an extraordinary toll on global health and the economy (1). COVID-19 patients include mild, moderate, even severe and critical COVID-19 cases (2, 3). Dysregulated host immune responses to SARS-CoV-2 infection may contribute to viral persistence, causing an amplified inflammatory response and subsequent lung injury and multi-organ damages, placing patients at a high risk of severe/critical COVID-19 (4).

The robust expansion of effector CD8^+^ T cells in vivo is an effective prerequisite for controlling acute viral infection. However, clinical research on circulating immune cells and lung sampling studies have documented a sharp decrease of CD8^+^ T cells in severe/critical cases (5, 6). Recent studies suggested that CD8^+^ T cells in severe or critically ill COVID-19 patients exhibited impaired effector function, limited proliferation and expansion compared to those in mild or moderate cases, despite high expression of some proliferation genes (7, 8), but the underlying mechanism is still poorly understood.

During virus infection, pathogen-specific CD8^+^ T cells are activated, proliferated and expanded to against virus infection, and cellular metabolism and mitochondrial function are the main determinants of this process (9, 10). CD8^+^ T cell activation is accompanied by extensive metabolic reprogramming with increased expression of nutrient transporter proteins and enzymes involved in glycolysis. In addition, effector CD8^+^ T cells rely on enormous energy generated by oxidative phosphorylation and glycolysis to complete the cell cycle (11,12). The mitochondrial oxidative phosphorylation system, a key system for energy supply in eukaryotic cells, comprises five enzymatic complexes (mitochondrial complex I to V) and two mobile electron carriers. Mitochondrial complexes I (NADH-CoQ reductase), II (succinate-CoQ reductase), III (CoQH2-c reductase), IV (cytochrome c oxidase), and V (oligomycin-sensitive ATPase) catalyze ATP production through a series of redox reactions, and adequate energy supply is of vital importance for CD8^+^ T cell proliferation and expansion. Therefore, insufficient cellular metabolism and mitochondrial energy generation might be responsible for the impaired functions and limited expansion of CD8^+^ T cells in severe/critical COVID-19.

In this study, through single-cell RNA sequencing (scRNA-seq) in combination with TCR sequencing, a proliferating-depleted CD8^+^ T cell phenotype with cell cycle arrest and impaired mitochondrial function was observed in severe/critical COVID-19 patients. Our ex vivo experiments further suggested that SARS-CoV-2 ORF3a in epithelial cells induced the galectin-3 expression, and caused the inhibition of mitochondrial complex gene transcription and mitochondrial biogenesis of CD8^+^ T cells, which was responsible for the cell cycle arrest and low expansion of CD8^+^ T cells. Our findings improve the understanding of the immune dysregulation during severe SARS-CoV-2 infection and provide support to develop novel, feasible, and effective treatments for COVID-19 infection.

## Materials and methods

### Research sources

#### Cell lines and plasmids

HEK293T cell line and A549 cell line were obtained from the American Type Culture Collection (ATCC, Manassas, VA, USA). The pLVX-EF1alpha-SARS-CoV-2-orf3a-2xStrep-IRES-Puro plasmid was constructed by Nevan Krogan lab, and provided by Addgene, USA. Lentiviral particles were packed using the 2^nd^ generation packaging system. Packaging plasmid GAG and envelope plasmid VSV-G were obtained from Addgene, USA.

### The data processing of single-cell RNA-seq and TCR sequencing

#### Single Cell Filtering, Clustering and Dimensionality Reduction

Subsequent calculations on the downloaded count matrix were performed using the R package Seurat (https://satijalab.org/seurat/). The cells were filtered based on the standard that the number of expressed genes was <200 or >2500, the UMI count was <500, or the mitochondrial gene percentage was >0.2,. The batch effect across different samples is removed by the “IntegrateData” function. The filtered expression matrix was normalized using the ‘LogNormalize’ method with a scale factor of 10000. The top 2000 variable features were then identified by the ‘vst’ method carried out by the Seurat function ‘FindVariableFeatures’. In order to make the data obey the normal distribution, the expression values of all genes were transformed by z-score using the ‘ScaleData’ function. Next, the principal component analysis (PCA) was performed through the top 2000 variable genes, and the data was dimensionally reduced based on the first 15 PCs. The ‘FindClusters’ function was implemented for clustering analysis on the PCA-reduced data subsequently, and the resolution was set to 1. Finally, the uniform manifold approximation and projection (UMAP) analysis was calculated on the top 15 principal components for visualization.

#### Cluster annotation

The initial cell cluster annotation was done based on canonical cell markers as follows: B cells, CD79A; CD4^+^ T cells, CD3E, CD4; CD8^+^ T cells, CD3E, NKG7, CD8A, and CD8B; NK cells, FCGR3A, NCAM1, and NKG7; Myeloid cells, ITGAM, CD68, FCGR3A and FCGR3B; and Epithlial cells: KRT18. To verify the validity of the cell-type assignment, the SingleR algorithm (https://github.com/dviraran/SingleR) was also performed independently.

#### Epithelial cells and CD8^+^ T cells re-integration

Epithelial cells and CD8^+^ T cells were re-integrated and re-clustered, respectively, with Seurat package. Specifically, these cellular data were all dimensionally reduced using PCA and UMAP based on the top 15 principal components with resolution parameter set to 0.8.

#### TCR V(D)J analysis

The scRepertoire package (v1.0.2) was used for single-cell immune repertoire data analysis. Combined with seruat package, the clonotype distribution of clusters defined by scRNA-seq data was displayed.

#### Trajectory inference and pseudotime

Monocle 3 (https://github.com/cole-trapnell-lab/monocle-release/), a pseudo-temporal inference algorithm, was utilized to reconstruct the cell differentiation trajectory. The CD8^+^ T cells from patients with severe/critical COVID-19 were selected for trajectory inference, and different branches in the cell trajectory may distinguish different cell states. Combined with the relevant biological background, we determined the “root” starting point, and eventually judged the order of cell population differentiation. In addition, Monocle 2 was used to further explore the cluster and lineage relationships between cluster 3 and cluster 10, the two specific cell clusters in CD8^+^ T cells.

#### Cell-cycle analysis

Cell cycle distribution of T cells was determined using the R package reCAT (https://github.com/tinglab/reCAT). The pipeline was applied with default and recommended parameters. We also use the ‘CellCycleScoring’ function in the Seurat package to evaluate the cell cycle and make cell cycle diagrams.

#### Gene set enrichment analysis

Gene set enrichment analysis (GSEA) was performed by the R package fgsea with C5, the Ontology Gene Sets, from the database Msigdb (https://www.gsea-msigdb.org/gsea/msigdb). The fgsea result was initially filtered based on the criteria that adjust p-value <0.05. For certain analyses, the downregulated mitochondria-related pathways from the GSEA table were selected according to the normalized enrichment score < 0, and the comparison results of related clusters were visualized via ggplot2 (v3.4.1).

#### Cellular communication

CellChat v0.0.2 (https://github.com/sqjin/CellChat) was used to assess cell-cell interactions based on the expression of known L–R pairs in different cell types. In particular, more than 100 related ligand-receptor pairs were added according to the official guidance from CellChat, for further analysis of the galectin pathways of concern. The results were visualized and displayed by related functions in the CellChat package.

### In vitro experiments

#### Generation of stable cell lines expressing SARS-CoV-2 ORF3a

Lentiviral packaging and amplification were performed for constructing stable cell lines using the 293T packaging cells. 293T cells were seeded in 6-well plates and reached 60-70% cell fusion rate next day. 293T cells were transfected with premixed plasmids and PEI (MW25000Da, Sigma, USA) and incubated for 72 h. The supernatants containing lentiviral particles were collected, and filtered by 0.22 μm filters. Lentiviral particles infected A549 cells with polybrene (Sigma, USA) at a final concentration of 8ug/ml. The A549 cells expressed SARS-CoV-2 ORF3a were selected with puromycin, and ORF3a expression in A549 was verified by western blot assay.

#### RNA extraction and Real-time PCR

Total RNA was extracted from cell samples using RNAsimple Total RNA Kit (Tiangen, CHN) according to the manufacturer’s instructions. The extracted RNA was then reverse transcribed into cDNA using ABScript II RT Mix (ABclonal, CHN). Real-time PCR was performed with 2×Universal SYBR Green kit (ABclonal, CHN). The fluorescence signal was collected by CFX96 PCR instrument (Bio-Rad, USA). Each sample was run in triplicate. The results were analyzed and examined as the relative mRNA levels based on cycle threshold (CT) values, which were converted to fold changes. The primers were designed using Primer-blast, NCBI (www.ncbi.nlm.nih.gov).

#### Western blot

The stable cell line was collected and lysed with RIPA lysis buffer (Servicebio, CHN) for 20 min on ice. Protein concentration was determined by a BCA protein concentration assay kit (Biosharp, CHN). Equal amounts of protein (2 μg/μL) were loaded on 10% gels and separated by SDS-PAGE, then electrophoretically transferred to PVDF membranes. Non-specific spots on PVDF membranes were blocked with 5% non-fat milk in PBST for 1 h. Membranes were incubated with primary antibodies (β-tubulin, Strep II-Tag, Abclonal, CHN) overnight at 4°C. HRP-coupled goat anti-rabbit IgG secondary antibodywas incubated for 1 h at RT. The protein bands were visualized using an ECL kit (Meilunbio, CHN) and imaged on ChemiDoc XRS+ (Bio-Rad, USA).

#### Isolation of CD8^+^ T cells and co-culture

The 24-well plates were coated with CD3 antibody (OKT3, Invitrogen, USA) overnight. PBMCs were isolated from the peripheral blood of healthy donors with Human PBMC Isolate (TBD, CHN) according to the manufacturer’s instructions. CD8^+^ T cells were then isolated by positive selection using magnetic beads coated with CD8 antibodies (BD, USA) and placed into the Transwell lower chamber in the presence of CD28 antibody (Invitrogen, USA) and IL-2 (Procell, CHN). A549 or A549-ORF3a cells were seeded in Transwell inserts (Corning, USA). For certain experiments, cells were treated with the galectin-3 inhibitor TD-139 (Selleck, USA) at a concentration of 15 μM. The co-culture system was incubated for 48 h at 5% CO2, 37°C.

#### Phosphokinase Array

Phosphoprotein expression of CD8^+^ T cells was detected using a human phospho-kinase array kit (ARY003C, R&D Systems, USA) according to the manufacturer’s protocol. Briefly, cell lysates were prepared and incubated on membranes containing human phosphokinase antibodies overnight at 4°C. Membranes were then incubated with detection antibody cocktail. After washing, HRP-conjugated secondary antibody cocktail was incubated on membranes. Chemi Reagent Mix was then incubated on membranes, and dot blots were visualized via chemiluminescence (Bio-rad, USA). The spot signals were quantified using Fiji software (https://imagej.net/Fiji).

#### Immunofluorescence staining and confocal laser scanning microscope analysis

CD8^+^ T cells were seeded to climbing slides at a density of 10^6^ cells/well. After fixation with 4% paraformaldehyde for 15 min, the cells were permeated with 0.3% Triton X-100 for 20 min, and then blocked for 1 h. NRF1 antibody (Abcam, USA) was used as the primary antibody and incubated overnight at 4°C. Rhodamine (TRITC) conjugated goat anti-rabbit IgG (H+L) were incubated for 1 h at room temperature. Then the slides were counterstained for nuclei and sealed with a DAPI-containing antifade mounting medium. NRF1 was visualized using a laser scanning confocal microscope (Olympus FV3000, JPN) with an excitation wavelength of 561 nm and a 100× silicone oil lens. Relative NRF-1 nuclear/cytosolic fluorescence ratios were quantified using ImageJ with the Fiji image processing package.

### Statistical analysis

For single-cell RNA-seq data analysis, R (v4.0.2) was utilized for informatics analysis and graph processing. All of the experiments were repeated at least three times, independently, with similar results. Statistical analysis was carried out using the t-test for two groups (GraphPad Prism 7). The date was considered statistically significant when *P ≤ 0.05, **P ≤ 0.01, ***P ≤ 0.001, ****P ≤ 0.001; ns stands for not significant.

## Results

### A dysfunctional CD8^+^ T cell subpopulation expressing proliferative signatures emerges in severe/critical COVID-19 patients

Ever since SARS-CoV-2 first appeared, researchers have been trying to understand how the immune system works during each stage of COVID-19. In this study, we firstly analyzed scRNA-seq data to measure the T cell responses against this virus and aimed to find out what caused the COVID-19 disease to develop into a severe or even critical condition.

Uniform Manifold Approximation and Projection (UMAP) was used to visualize the clusters of all the cells identified by a shared nearest neighbor (SNN) modularity optimization-based clustering algorithm implemented in Seurat v4. 27 distinct clusters were shown by clustering analysis, covering diverse cell types in the respiratory system (Figure 1A). Major groups of immune cells in BALF, namely macrophage, neutrophil, T, B, NK, and epithelial cells were identified by combining specific gene expression signatures with the SingleR method (14). The expression levels of major signatures to annotate these subpopulations were shown in Figure 1B.

**Figure 1.**
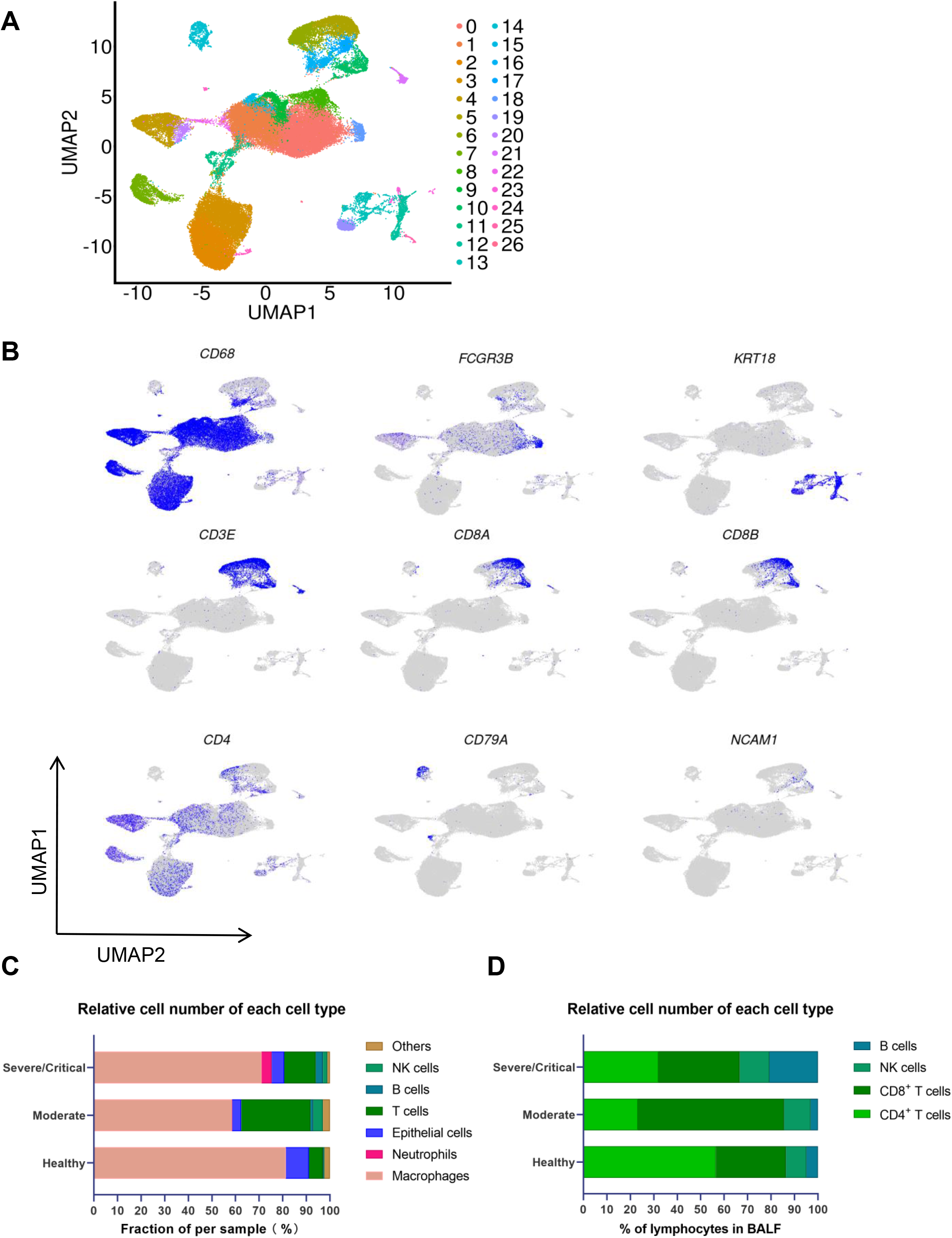
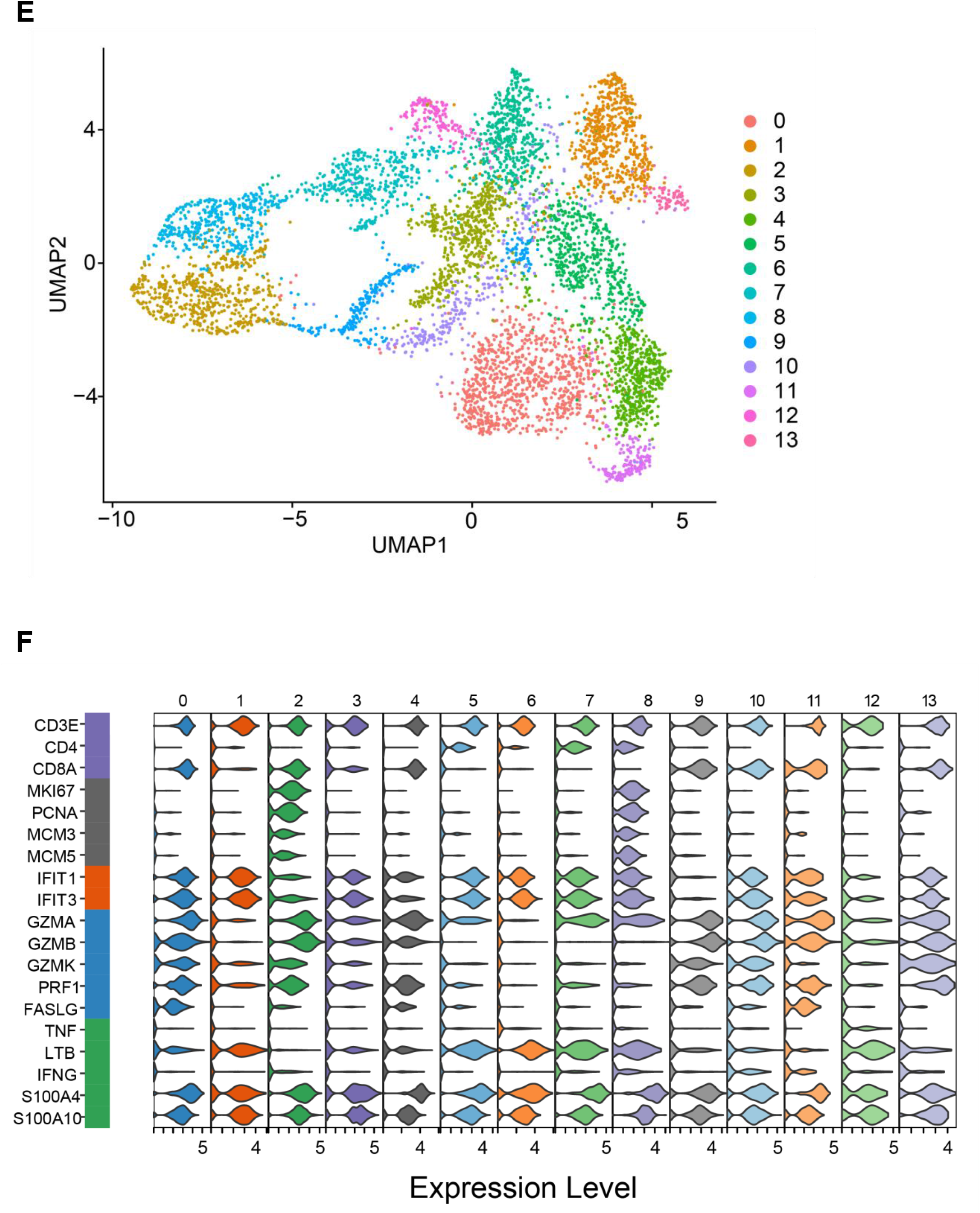
Two distinct T cell subpopulations are associated with COVID-19 severity. (A) Overview of the cell clusters in the integrated single-cell transcriptomes of BALF cells derived from COVID-19 patients and healthy controls. UMAP of 27 cell clusters among healthy controls, moderate COVID-19 patients and severe/critical COVID-19 patients were displayed. (B) Marker genes used to identify major cell types were specifically expressed in the corresponding clusters. (C) The bar plot compares the proportion of major BALF cell types in healthy controls, patients with moderate and severe/critical COVID-19. (D) The bar plot showed the percentage of lymphocyte clusters in healthy controls, patients with moderate and severe/critical COVID-19. (E) The UMAP of 14 heterogeneous clusters of T cells isolated from BALF of COVID-19 patients. (F) Violin plots showed normalized expression level of representative phenotypic (CD3E, CD4, CD8A), proliferation (MKI67, PCNA, MCM3, MCM5), IFN-stimulated (IFIT1, IFIT3), cytotoxic/functional (GZMA, GZMB, GZMK, PRF1, FASLG) and inflammatory marker transcripts (TNF, LTB, IFNG, S100A4, S100A10) respectively in T cells from BALF of COVID-19 patients.

We then analyzed the proportions of each cell type to find the groups that played a key role in the progression of COVID-19. In the BALF of moderate patients, the proportion of T cells was 29.33%, much higher than that of severe/critical COVID-19 patients (13.37%) and healthy people (5.95%) (Figure 1C). Significant T cell expansion occurred in moderate COVID-19 patients, which is consistent with previous related reports (6, 15). However, compared with moderate patients, the percentage of T cells in severe/critical patients was reduced to a certain extent. For the composition of lymphocytes, CD4^+^ T cells in moderate patients accounted for 23.24%, and CD8^+^ T cells accounted for 62.36%, while in severe/critical patients, CD4^+^ T cells and CD8^+^ T cells respectively accounted for 32.04% and 34.63% (Figure 1D). Patients with severe/critical COVID-19 had more CD4^+^ T cells, and their CD8^+^ T cells were drastically decreased as opposed to moderate patients.

To further explore the characteristic changes in severe patients, we turned to subset and re-clustered the T cell data by UMAP. For the follow-up analysis, the cells of healthy people were excluded. To prevent the mixing of other cell types, the selected T cell data were filtered again according to correspondingly typical markers. Ultimately, other types of cells like myeloid cells and epithelial cells were basically excluded and the re-screened T cell population was divided into 14 distinct clusters (Figure 1E). Cluster 2 and cluster 8 exclusively displayed high levels of proliferation markers (MKI67, PCNA, MCM3, and MCM5). These two subpopulations emerged at the stage of severe/critical disease, whereas they were derived from CD8^+^ T and CD4^+^ T cells, respectively. It is noted that cluster 2 seemed to be a proliferative subset under this circumstance with abundant cell numbers, while in fact, the proportion of CD8^+^ T cells decreased in patients with severe/critical COVID-19 (Figure 1D). On the other hand, transcript levels of functional genes like FASLG was also decreased in cluster 2. Down-regulated inflammatory factors like LTB and TNF also revealed that the cellular function of cluster 2 has been somewhat impaired. In the meanwhile, cluster 2 also displayed high cytotoxic signatures (GZMB and PRF1), intermediate levels of inflammatory proteins (S100A4 and S100A10), and partially remained IFN inducibility, suggesting a disordered status influenced by pathologic lung condition (Figure 1F).

### Critical CD8^+^ T cell subpopulations have cell cycle arrest and are correlated with the disease process of COVID-19

Now single-cell analysis revealed a group of abnormal CD8^+^ T cells originating from severe/critical patients and expressing proliferative genes, which conflicts with the current situation of CD8^+^ T cell reduction. In order to further interrogate the critical role of CD8^+^ T cells found above in severe/critical COVID-19 patients, we then selected all CD8^+^ T cells from the total cells of BALF. Characteristic signatures of CD8^+^ T cells and others were identified, and the varying degree of expression of which in specific cell clusters was visualized. Finally, 3870 CD8^+^ T cells were isolated to perform subsequent analyses. We observed that these CD8^+^ T cells highly expressed both T cell markers and CD8^+^ T cell markers, and they had almost no expressions of the CD4^+^ T cell, myeloid cell, or epithelial cell markers. Thus, we identified that CD4^+^ T cells, myeloid cells, and epithelial cells were almost excluded from our analyses, and obtained relatively pure CD8^+^ T cell data.

These CD8^+^ T cells were then clustered into 13 subpopulations and were clearly differentiated according to their sample origin (Figure 2A, B). Proportions of CD8^+^ T cells in different disease stages for each cell cluster were shown in Figure 2B. These clusters showed a distinguishing pattern in the low-dimensional distribution, with cluster 6 consisting only of cells of healthy origin distinct from other clusters of cells nearly all from COVID-19 patients, which revealed that CD8^+^ T cells of COVID-19 patients and healthy people have obvious heterogeneity. The identical high expression of proliferative genes showed that C3 (stand for cluster 3) and C10 (stand for cluster 10) were two subsets of cluster 2 above, and also the key cell populations of our interest (data not shown). In order to examine the extent of their expansion, the clonotypes were further analyzed. Consistent with the decline in the proportion of CD8^+^ T cells, the clonotype size of CD8^+^ T cells derived from severe/critical patients in C3 and C10 was lower than that of corresponding moderate patients (Figure 2C), indicating that cell cloning is actually hindered.

**Figure 2.**
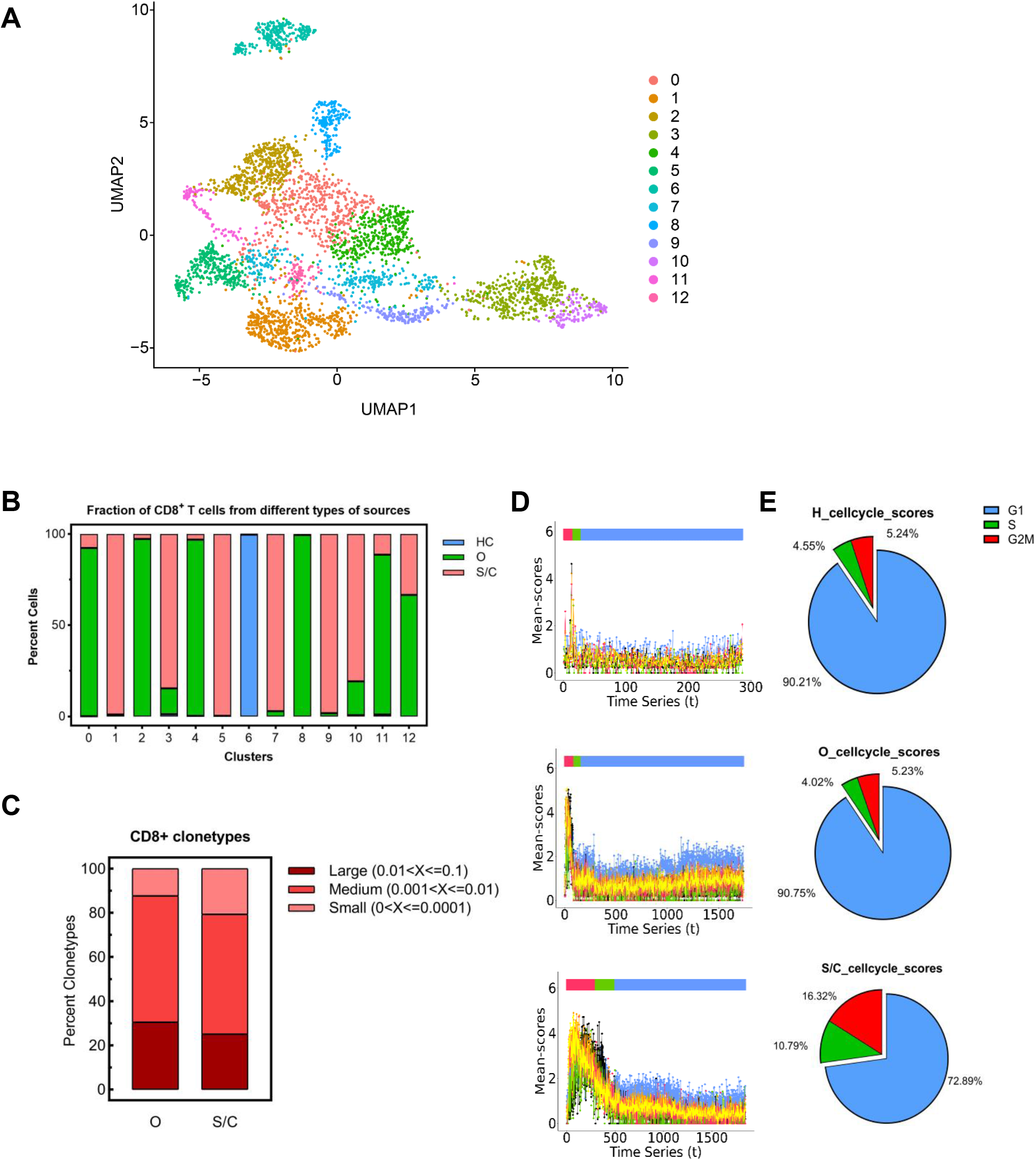
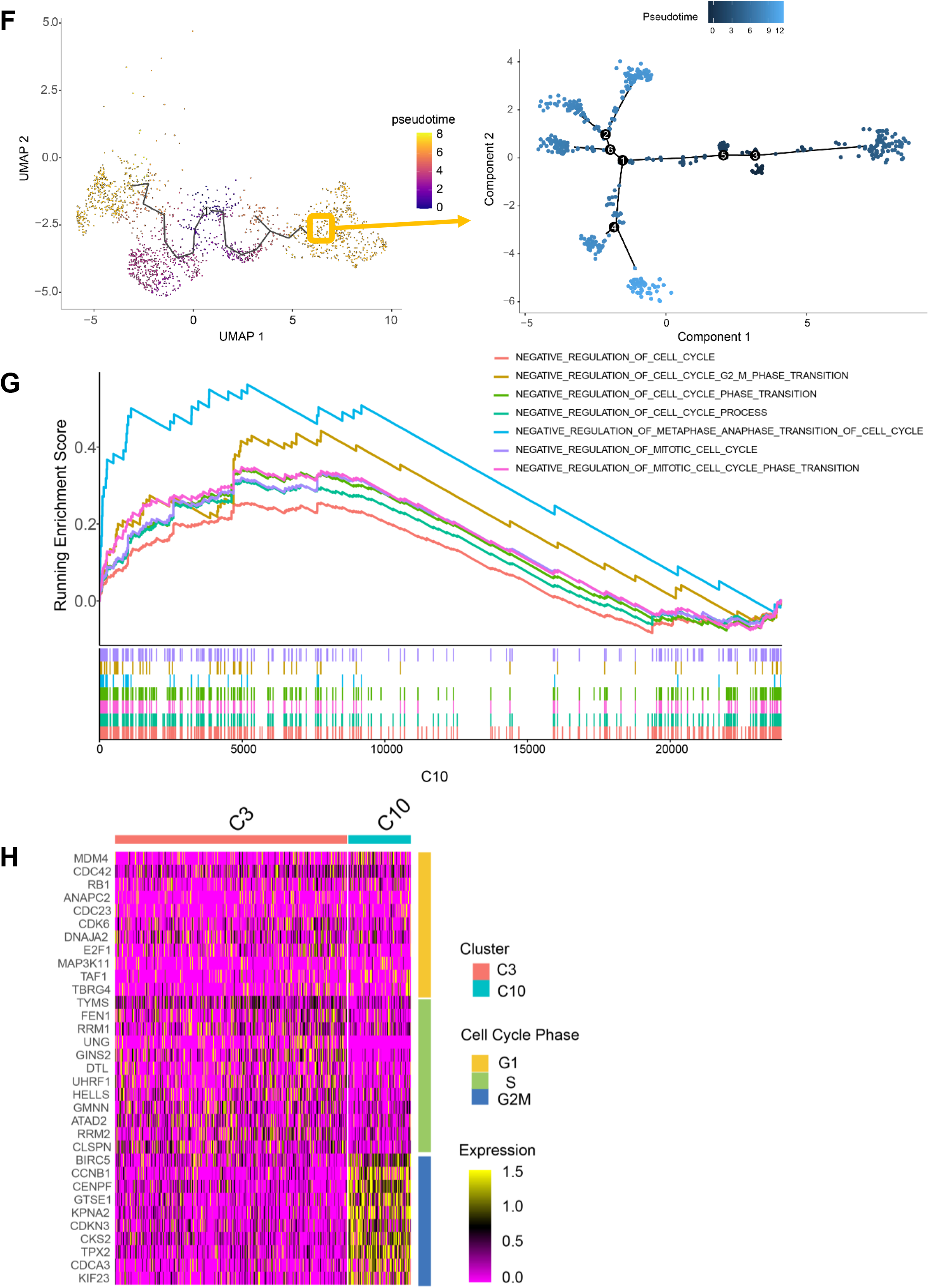
Cell cycle arrest occurred in certain CD8^+^ T cell subpopulations of severe/critical COVID-19 patients. (A) UMAP embedding of all CD8^+^ T cells colored by unsupervised clustering. (B) Percentage of CD8^+^ T cells across healthy controls, moderate, and severe/critical COVID-19 patients in individual clusters. HC means healthy controls, O means moderate COVID-19 patients, and S/C means severe or critical COVID-19 patients. (C) Bar plots showed the percentage of clonotypes in specified CD8^+^ T cell cluster from patients with moderate and severe/critical COVID-19. The clonotypes are categorized as Large (0.01<X<=0.1), Medium (0.001<X<=0.01) and Small (0<X<=0.0001) based on their relative abundance. (D) reCAT reconstructs cell cycle time-series and predicts cell cycle stages along the time-series. The presentation of different cell cycle phases of healthy controls (upper panel), moderate patients (middle panel) and severe/critical patients (lower panel). (E) The corresponding pie charts showed the proportion of CD8^+^ T cells at different cell cycle stages. (F) Pseudo-time trajectory projected onto a UMAP of selected CD8^+^ T cells. Pseudo-time values are color coded. Numbers in circles indicate inferred differentiation paths. The enlarged box is the timing diagram of C3 and C10 (located at the bottom of the picture). C3, 10: two CD8^+^ T cell groups isolated from severe/critical COVID-19 patients. (G) Enrichment plots for pathways identified by GSEA. C10 positively correlated with the negative regulation of cell cycle relative signaling pathway in the Molecular Signatures Database (MSigDB). (H) Heatmap showing average expression of cell cycle phases gene signatures (rows) in C3 and C10 (columns). The color scale shows the expression level of selected transcripts in each cell.

To further investigate the contradiction behind this, we analyzed the cell cycle of these CD8^+^ T cells with reCAT (16). The cell cycle is known to have wide-ranging effects on cellular physiology and can modulate both differentiation and gene expression profiles. Proliferation depends on the smooth completion of four distinct phases of the cell cycle — G0/G1, S, G2, and M (17). As shown in Figure 2D, severe/critical COVID-19 patients had significantly more CD8^+^ T cells in G2/M phase than that in moderate patients and healthy people. Among CD8^+^ T cells of patients with severe/critical COVID-19, 10.79% were in S phase and 16.32% were in G2/M phase. In contrast, the cell cycle distributions of moderate patients and healthy people were similar, with 4.02%/4.55% in S phase and 5.23%/5.24% in G2/M phase (Figures 2D, E). These results indicated that the CD8^+^ T cells sourced from severe/critical COVID-19 patients may have cell cycle arrest.

MKI67, which is associated with and may be necessary for cellular proliferation, was only expressed in C3 and C10 (data not shown). In the meanwhile, we also found MCM4 and MCM5, which are essential for the initiation of eukaryotic genome replication, were exclusively highly expressed in C3 (data not shown). Pseudo-time analysis revealed a differentiation trajectory of CD8^+^ T cells sourced from severe/critical COVID-19 patients, which showed C10 was at the rear end of C3 (Figure 2F). Collectively, these results raised the possibility that C10 may be developed from C3.

GSEA analysis showed that C10 had numerous up-regulated signaling pathways about negative regulation of cell cycle (Figure 2G). To be more precise, we next checked the expression of cell cycle-related genes, to find that C10 mainly expressed G2/M phase genes like BIRC5, CCNB1, CENPF, GTSE1, KPNA2, CDKN3, and CKS2, while C3 had a universal expression pattern with no obvious cell phase preference (Figure 2H). Taken together, C10 may be a cluster of cells that could not complete amplification due to cell cycle arrest at G2/M phase, most of which came from severe patients, suggesting a potential reason why CD8^+^ T clonotypes were not enlarged in severe cases.

### Impairment of mitochondrial function in the cell-cycle-arrest cluster

T cell proliferation and activation require ATP produced by mitochondrial activity (18). To further explore the cause behind cell cycle arrest in C10, we picked out C3 and C10 cell subpopulations from severe/critical COVID-19 patients and observed their mitochondrial respiratory chain complex-related gene expression. All of these respiratory chain complex genes are nuclear-coding genes. Mitochondrial complex-related genes were mostly expressed in C3, and downregulated in C10 (Figure 3A). Representative genes like NDUFA4, NDUFB2, NDUFB11, NUDFS5 (encoding mitochondrial respiratory chain complex I), UQCRB, UQCRQ, UQCRH UQCR10 (encoding complex III), COX4I1, COX6A1, COX6C and COX7A2 (encoding complex IV), had relatively high expression levels in C3 but almost no expression in C10 (Figure 3A). We next collected mitochondria-related signaling pathways from the MSigDB database and performed GSEA analysis on C3 and C10, separately.

**Figure 3.**
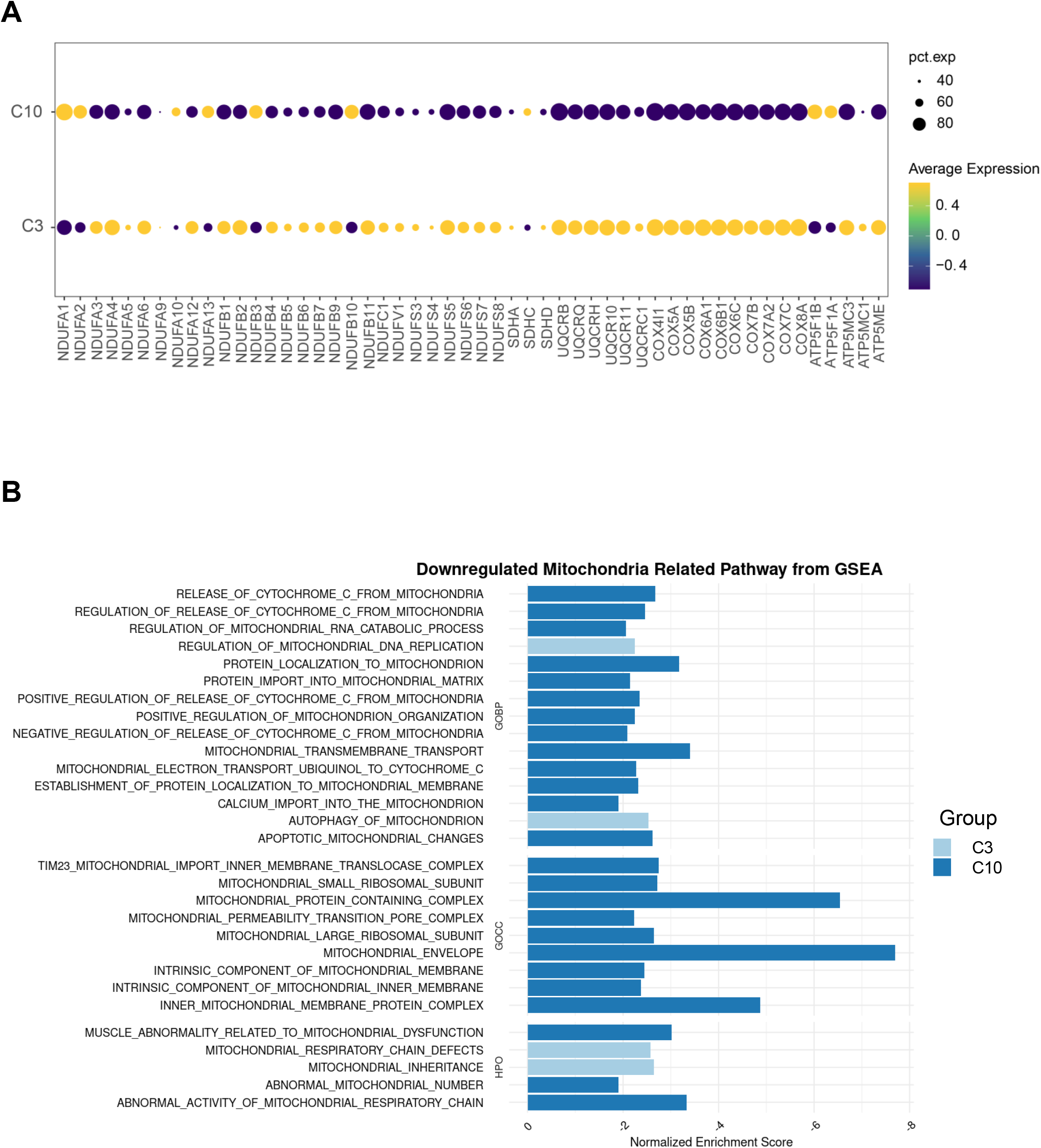
Impaired mitochondrial function in CD8+ T cells from severe/critical COVID-19 patients. (A) The expression of mitochondrial complex-related genes in different cell subsets (C3 and C10) is depicted by dot plots. Pct. Exp.=percentage of cells expressing the gene. Color scale shows the average expression level of mitochondrial complex-related genes. (B) Gene Set Enrichment Analysis (GSEA) for specific CD8^+^ T cells in sever/critical COVID-19 patients on downregulated mitochondria related pathways. Column chart showing the normalized enrichment scores for the mitochondria related pathways derived from BP, CC and HPO that are significantly downregulated in C3 or C10.

Enrichment results of C10 showed plenty of downregulation of mitochondrial-related pathways, while enriched terms in C3 had little relationship with mitochondrial function (Figure 3B). This strong decrease of respiratory chain-related genes and mitochondrial-related signaling pathways revealed that the mitochondrial biogenesis of C10 was suppressed, and its paramount function — supply energy for life activities — was impaired.

### Significant galectin-associated interactions between lung epithelial and abnormal CD8^+^ T cells in severe/critical COVID-19 patients

The function and phenotype of immune cells are largely influenced by the environment, and the target cells of immune responses are often involved in reshaping the immune environment and regulating immune cells through different ways such as contact or secretion (19). We re-clustered 6 subclusters of lung epithelial of BALF to further dissect their heterogeneity and explore its effect on cluster 10 of CD8^+^ T cells. In case it mixed with other types of cells, the data of epithelial cells was filtered again according to correspondingly typical markers. Ultimately, the re-screened epithelial cell population was divided into 14 subpopulations and basically excluded other types of cells like myeloid cells and lymphocytes (Figure 4A). There were 3338 epithelial cells in total, of which 1456 cells were from healthy people, 211 cells were from moderate cases, and 1671 cells were from severe/critical cases, and their distribution in each cell population was graphed in Figure 4B. Patients with severe/critical illness experienced more SARS-CoV-2 invasion and replication in their respiratory tract, and thus more shed epithelial cells can be obtained in the alveolar lavage fluid (20), which could explain the difference in the number of epithelial cells from different sample sources.

**Figure 4.**
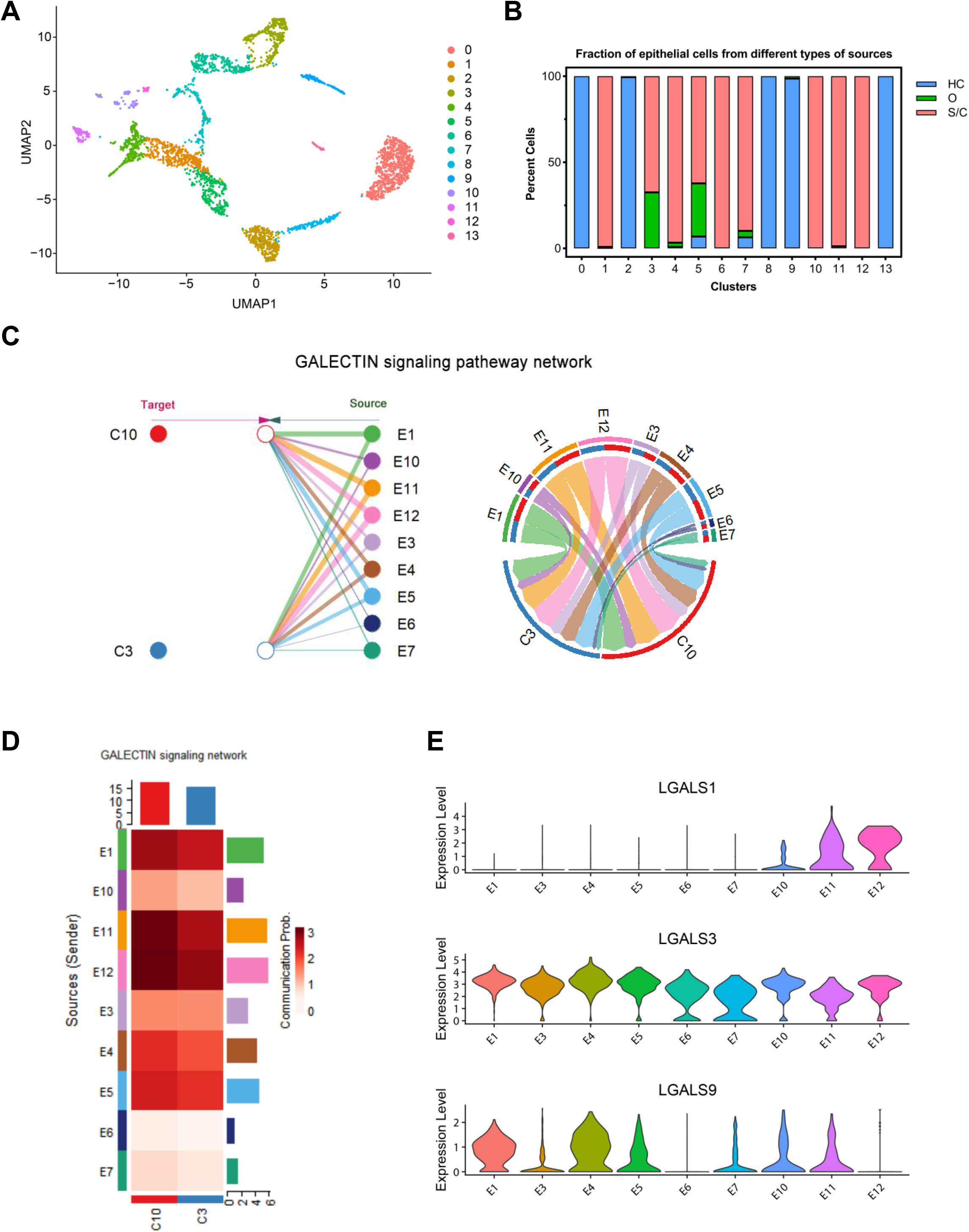
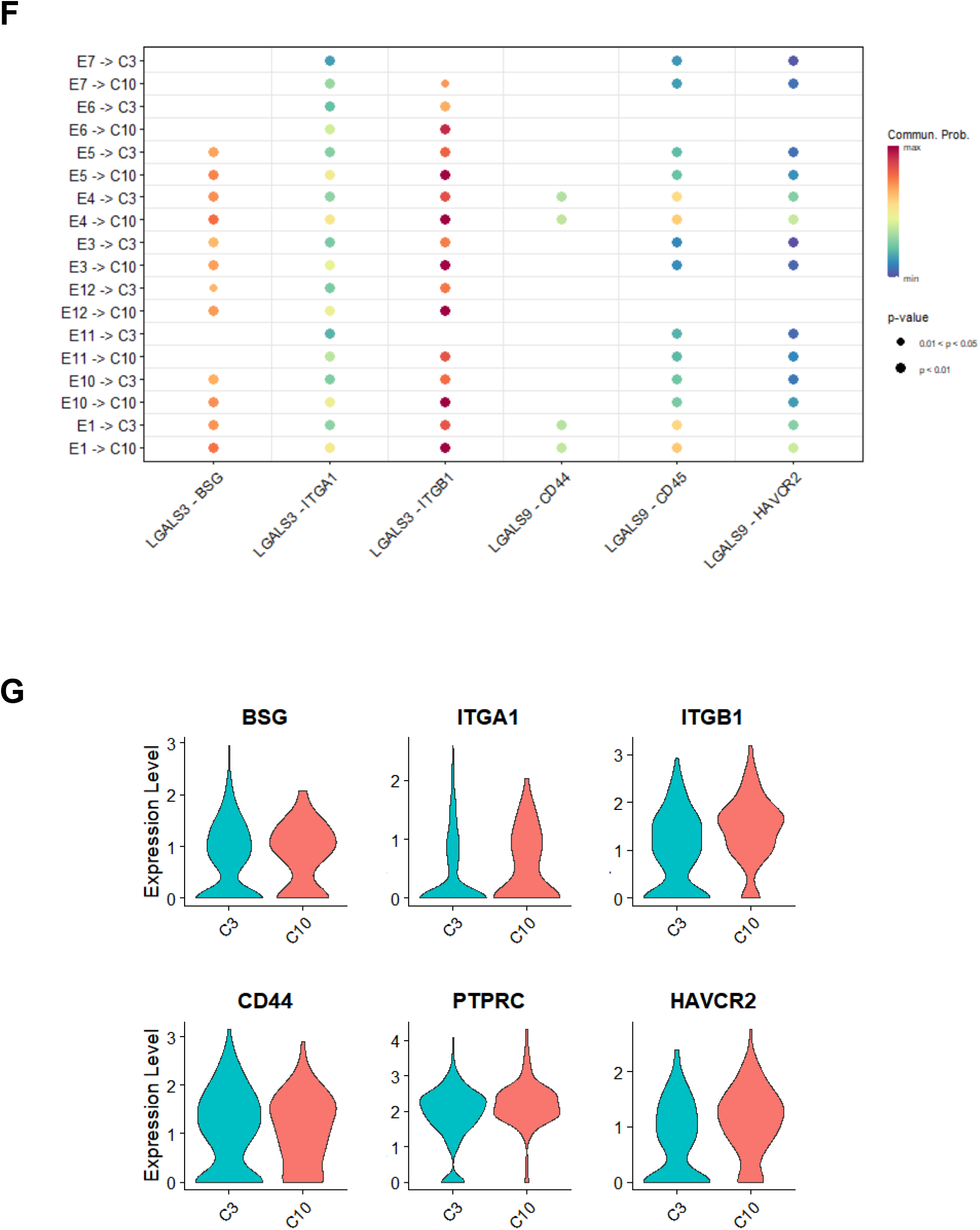
Galectin signaling pathways may be involved in mitochondrial dysfunction of CD8^+^ T cells in severe/critical COVID-19 patients. (A) The UMAP presentation of 14 heterogeneous clusters of epithelial cells. (B) Percentage of epithelial cells across healthy controls (HC, blue), moderate (O, green) and severe/critical (S/C, red) COVID-19 patients in individual clusters. (C) Hierarchical plot (left) and chord diagram (right) show the inferred intercellular communication network (only the effect of epithelial cells on CD8^+^ T cells is represented here) for galectin signaling. E1, 3-7, 10-12: nine epithelial cell groups separated from severe/critical COVID-19 patients; C3, 10: two CD8^+^ T cell groups from severe/critical COVID-19 patients. (D) CellChat infers the strength of different cell groups as senders or receivers of signals during cellular communication. The color bar shows the strength of signals, the histograms with different colors indicate the total strength of different cell groups, and the x- and y-axes represent the signal receiver (CD8^+^ T cells) or sender (epithelial cells). (E) Violin plots comparing the normalized expression level of LGALS1, LAGLS3 and LGALS9 transcripts in various epithelial clusters of COVID-19 patients with severe/critical disease. (F) Bubble plot showing stronger effects of multiple galectin ligand-receptor pairs on C10 involved in cellular communication. (G) Violin plots exhibit the expression levels of relative receptor genes in C3 and C10.

We choosed lung epithelial cells basically derived from severe/critical COVID-19 patients to conduct cell communication analysis with C3 and C10. R toolkit for single-cell data, CellChat (v1.5.0) was applied to analyze the signaling pathways that are possibly involved in epithelial-CD8^+^ T cell interactions. Galectin signaling pathway was found especially prominent among all the secreted signaling pathways (Figure 4C). Galectins are a family of fifteen β-galactoside-binding proteins, which play different roles in various biological processes, such as cell cycle regulation, apoptosis, and cytokine synthesis of effector helper and cytotoxic T cells. In general, galectins are known for their pro-adhesive potential and their negative effects on T cell proliferation and survival (21). However, the limitation is that the CellChat official database is insufficient to explore for it only contains three ligand-receptor pairs involved in the galectin signaling pathway. In order to investigate the role of galectin-related interactions between epithelial cells and CD8^+^ T cells from severe/critical COVID-19 patients more comprehensively, an extra ligand-receptor interaction list containing more than one hundred ligand-receptor pairs was supplemented according to official instructions.

For the convenience of description, we abbreviated the subpopulations derived from epithelial cells of severe/critical COVID-19 patients as E1, E3, E4, E5, E6, E7, E10, E11, and E12; as for the two key CD8^+^ T cell populations mentioned above were referred to as C3 and C10. The importance of each cell cluster in the galectin signaling pathway across all ligand-receptor interactions was computed by Cell Chat. E1, E4, and E5, the three subpopulations with a higher cell count, had a significantly obvious effect on C3 and C10 through the galectin signaling pathway (Figure 4D). Moreover, for each epithelial cell cluster, the likelihood or strength of their interaction with C10 was higher than that with C3 (Figures 4C, D).

The expression of several molecules of the galectin family that worked as ligands in epithelial cells is shown in Figure 4E. LGALS3 and LGALS9, encoding galectin-3 and galectin-9 respectively, were expressed highly in E1 and E4 (Figure 4E). In the ligand-receptor analysis of LGALS3 and LGALS9, it was found that there were six ligand-receptor interactions total, named LGALS3-BSG, LGALS3-ITGA1, LGALS3-ITGB1, LGALS9-CD44, LGALS9-CD45, and LGALS9-HAVCR2, showing E1 and E4 had a stronger effect on C10 than C3 (Figure 4F). Moreover, receptor mentioned above such as ITGA1, ITGB1, PTPRC, and HAVCR2 were up-regulated in C10 compared to C3 (Figure 4G). LGALS3 or LGALS9 binding to these receptors might cause negative regulation of CD8^+^ T proliferation and cell function.

### SARS-CoV-2 ORF3a induces upregulation of galectin-3 and downregulation of mitochondrial respiratory chain-related genes in A549 cells

We then verified and further explored the above findings through ex vivo experiments. ORF3a is an accessory protein encoded by SARS-COV-2, which not only facilities viral release, but also has been shown to affect a variety of physiological processes of host cells, including inducing apoptosis, blocking autolysosome formation, and triggering inflammatory responses (22, 23). Thus, we wondered if SARS-CoV-2 ORF3a (referred to as ORF3a hereafter) could cause the increased expression of galectins in epithelial cells, and mediate the downstream changes in CD8^+^ T cells.

To verify the assumption, A549 human lung epithelial cells were infected with SARS-CoV-2 ORF3a lentivirus, and a stable cell line (termed A549-3a) was generated after puromycin selection. The production of ORF3a proteins was subsequently detected and confirmed by western blot (Figure 5A). Next, the role of ORF3a in regulating proinflammatory cytokine production was evaluated. The results showed that Galectin-3, S100A14, IL-1β, and CCL2 mRNAs were induced upon ORF3a transfection into A549 cells (Figure 5B). Besides, ELISA confirmed the increase of galectin-3 in the cell culture supernatant (Figure 5C). These results were in line with the previous view that ORF3a causes inflammation, and confirmed our conjecture that galectin-3 can be induced by ORF3a.

**Figure 5.**
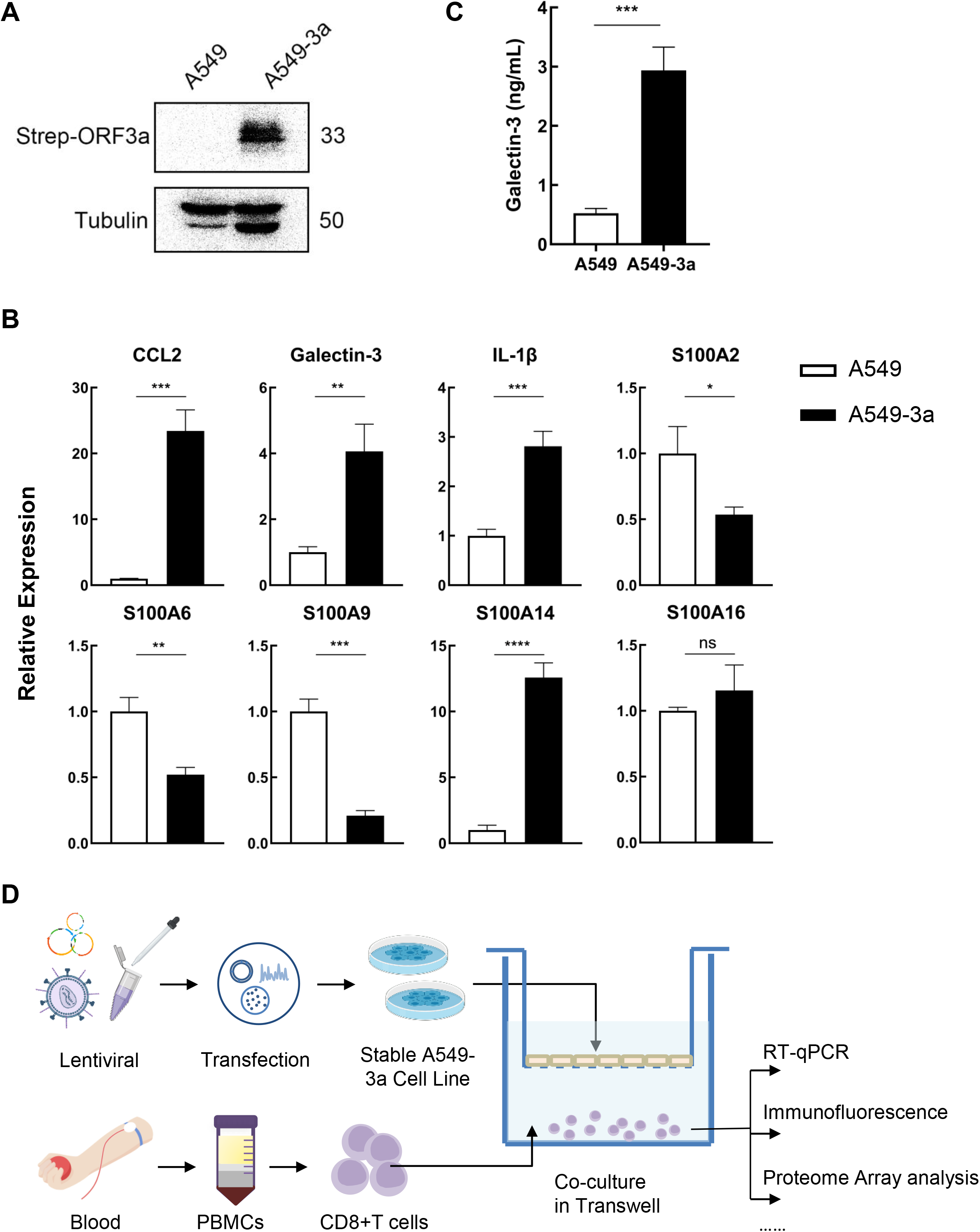
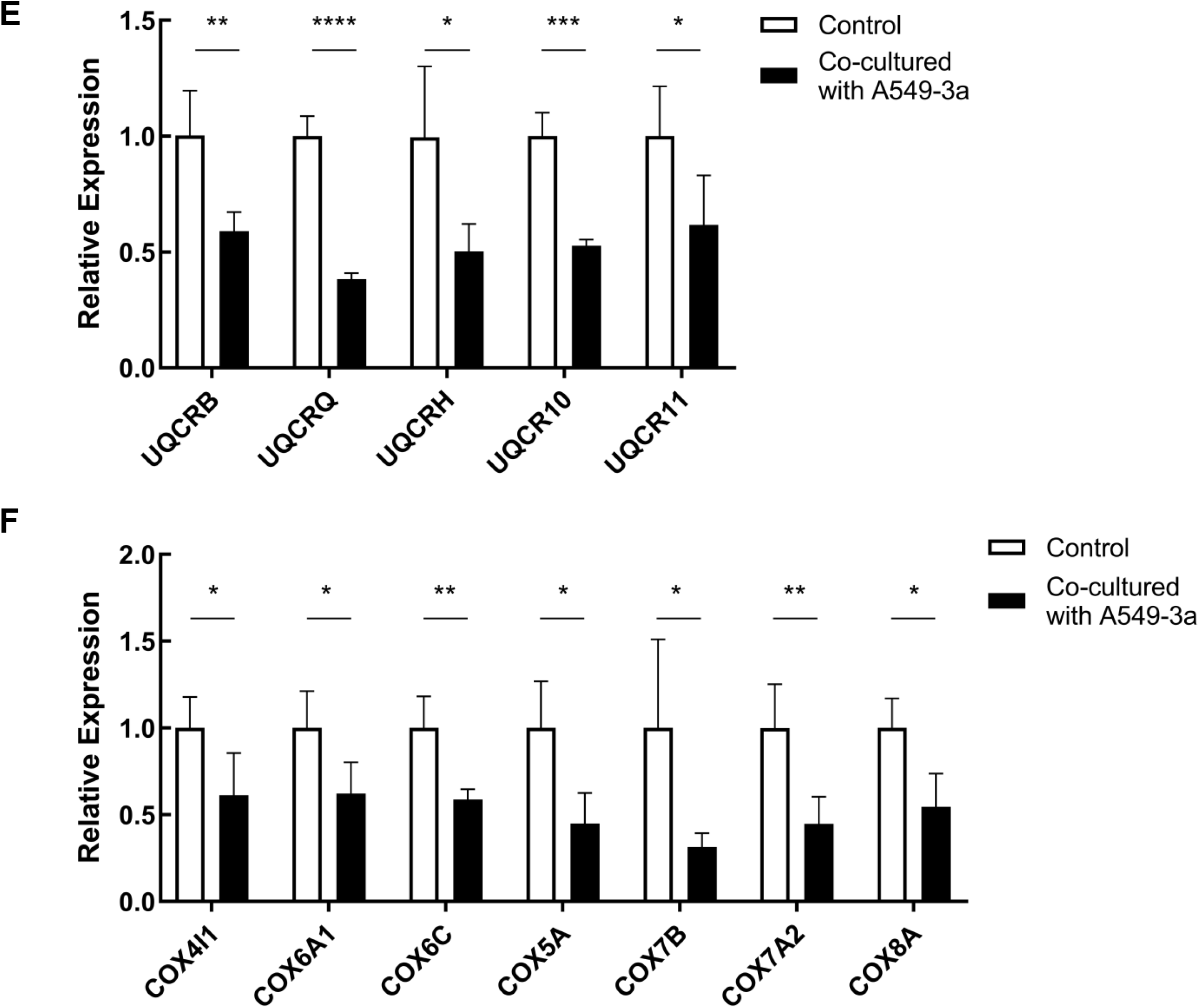
SARS-CoV-2 ORF3a induced galectin-3 expression in A549 cells and impaired mitochondrial biogenesis of CD8^+^ T cells. (A) Representative western blot images (with anti-Strep antibody) showing the expression of SARS-CoV-2 ORF3a in transfected A549 cells. A549 represents the control group, while A549-3a represents the treatment group transfected with SARS-CoV-2 ORF3a (B) Inflammatory S100 proteins, IL-1β, Galectin-3 and CCL2 mRNA levels in A549/A549-3a cells was assessed by Real-time-PCR. Relative gene expression was normalized against GAPDH. (C) Galectin-3 expression level in supernatant was detected by ELISA. (D) Experimental design to characterize the alternation of CD8^+^ T cells after co-culture with epithelial cells expressing ORF3a. CD8^+^ T cells were isolated from PBMC of healthy donor to co-culture with A549/A549-3a cells for 36 h, and then subjected to multiple assays. (E, F) The mRNA expression levels of ETC complexes III (E) and IV (F) in CD8^+^ T cells after co-culture with A549/A549-3a cells were determined by Real-time PCR. *P < 0.05; **P < 0.01; ***P < 0.001; ****P < 0.0001 with Student’s t-test (B) (C) or multiple t-test (E) (F).

To study the interaction between epithelial cells and CD8^+^ T cells in vitro, we designed the A549-3a/CD8^+^ T cell co-culture system and the experimental design is shown in Figure 5D. In brief, regular A549 or A549-3a cells were seeded on transwell inserts; CD8^+^T cells were isolated from peripheral blood of a healthy donor, co-cultured with the epithelial cells for 36 hours with anti-CD3/anti-CD28 stimulation, then subject to multiple detections. The qPCR results showed that after co-culture with A549 cells infected with ORF3a, the gene expression levels of mitochondrial complex III/IV of CD8^+^ T cells were significantly reduced (Figure 5E, F). Among them, the expression of UQCRQ, UQCR10, UQCRB, COX6C and COX7A2 decreased statistically significantly from the control group (p < 0.01). This suggested that epithelial cells infected with SARS-CoV-2 could downregulate mitochondrial complex-related gene expression and impair mitochondrial biogenesis in CD8^+^ T cells.

### Galectin-3 inhibits mitochondrial complex III/IV genes transcription by suppressing NRF-1

Our single-cell transcriptome analysis revealed that galectin-3-related ligand-receptor interactions between lung epithelial cell clusters and CD8^+^ T cell clusters were significantly enhanced in CD8^+^ T cell subsets with impaired mitochondrial function and cell cycle arrest. To investigate the role of galectin-3 in impaired mitochondrial biogenesis in COVID-19 patients, TD-139, a high-affinity inhibitor of galectin-3, was utilized in the A549-3a/CD8^+^ T cell co-culture system. As speculated, the transcriptional repression of COX genes and UQCR genes were differentially rescued with inhibitor treatment (Figure 6A).

**Figure 6.**
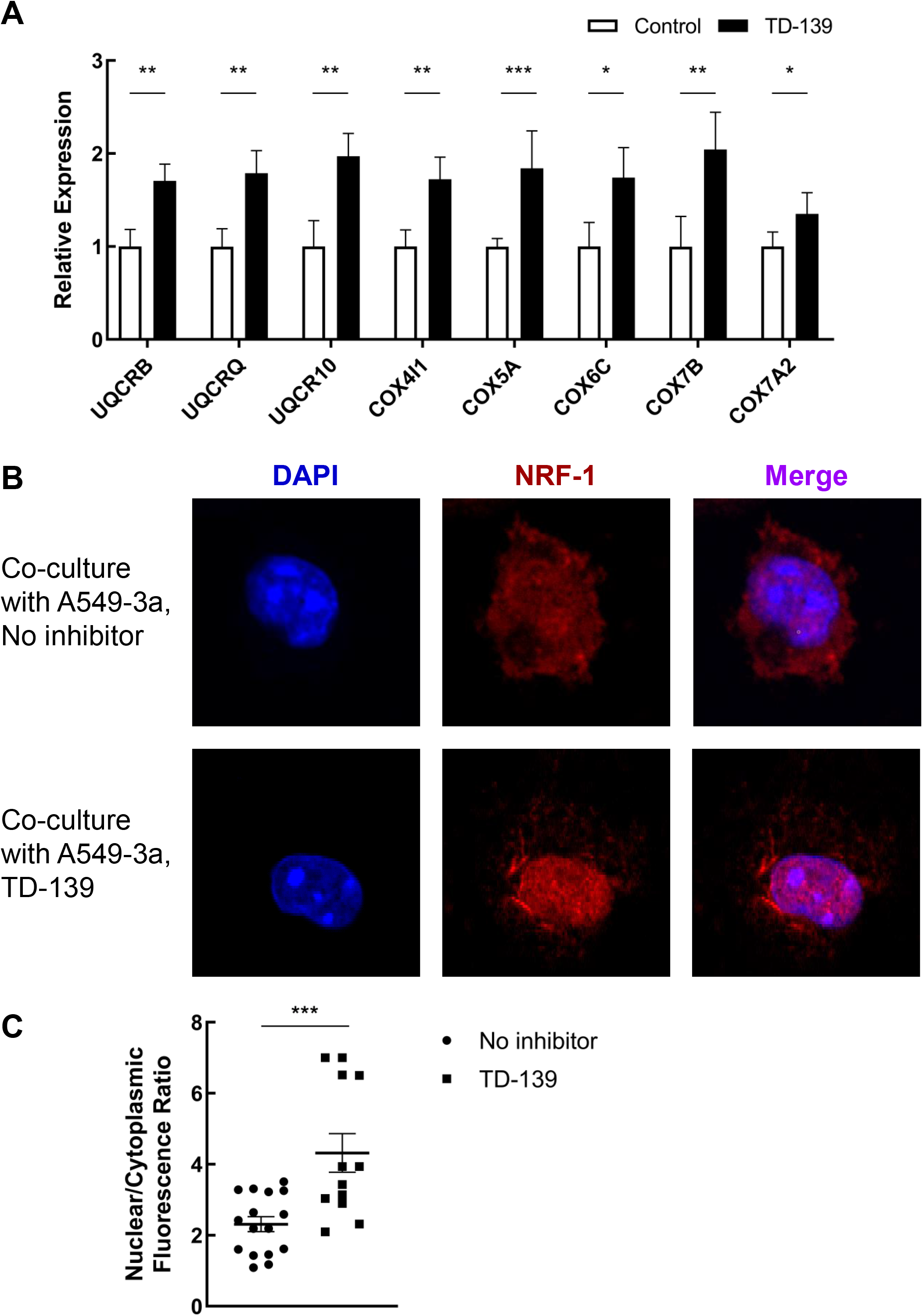
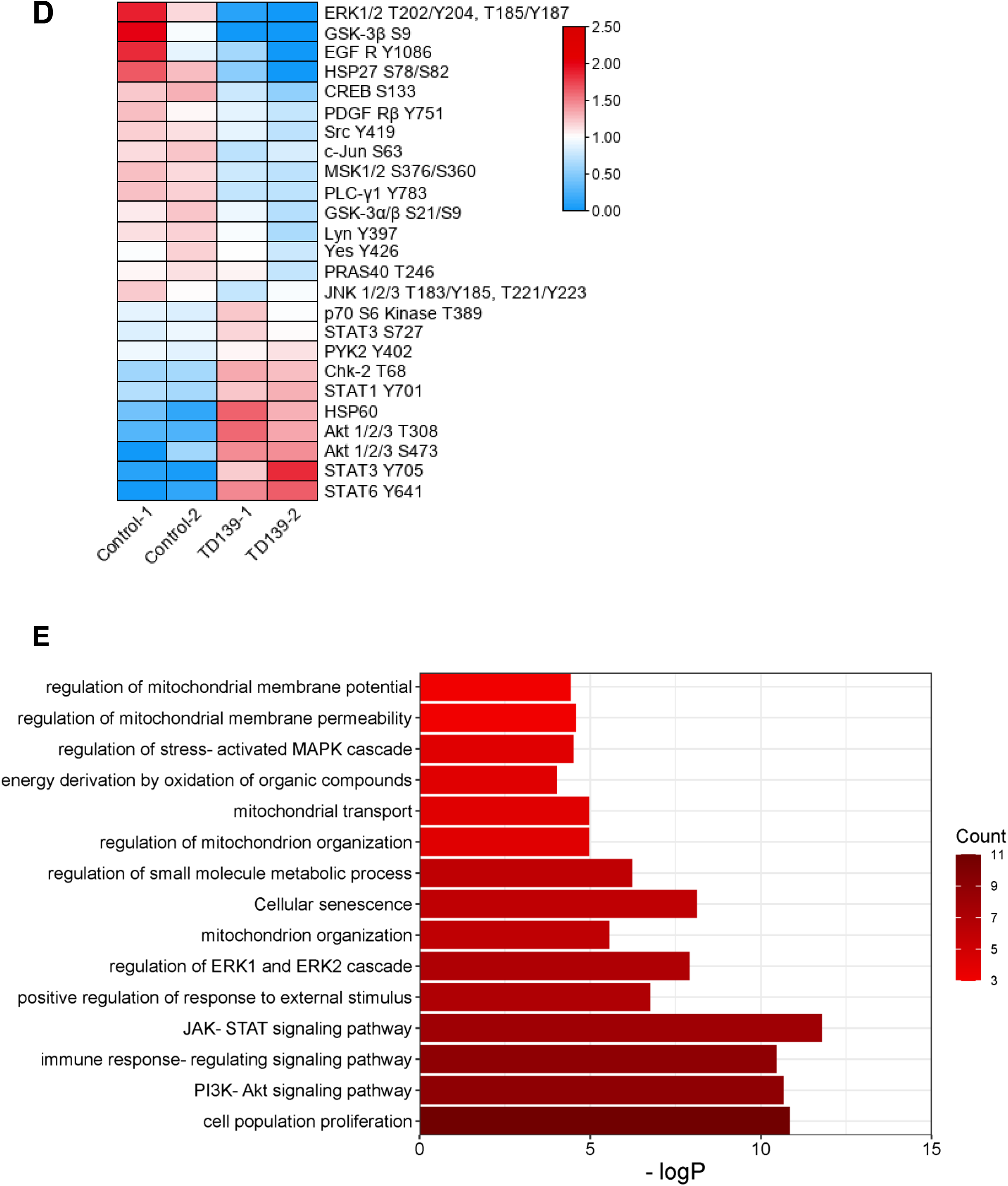
Galectin-3 inhibited ETC III/IV genes transcription by suppressing NRF-1 and was involved in ERK/Akt pathway. (A) The expression of ETC complexes III and IV genes in CD8+ T cells co-cultured with A549-3a cells. The TD-139 group was treated with galectin-3 inhibitor TD-139 for 48 h. (B) CD8^+^ T cells were examined by confocal microscopy for the localization and expression of NRF-1 (red). Nuclei were visualized using DAPI counterstain (blue). (C) Quantification of nuclear fluorescence intensities of NRF-1. (D) Phospho-Kinase production of CD8^+^ T cells co-cultured with A549-3a cells for 24 h in the presence or absence of TD-139 was compared using a commercial protein array kit. After quantification, 25 differentially expressed proteins were identified and visualized in heatmap. The color bar on the left represents the relative expression level. (E) Bar diagram of enrichment analysis of differentially expressed proteins involved in D. Color indicates the number of genes involved in specific signaling pathway. *P < 0.05; **P < 0.01; ***P < 0.001 with Student’s t-test (B) (C) or multiple t-test (E) (F).

In order to meet the energy and metabolic needs of cell proliferation, the number and function of mitochondria are finely regulated, which is largely accomplished at the transcriptional level (24). Previous studies showed that nuclear respiratory factor 1 (NRF1), a transcription factor for a wide range of genes involved in mitochondrial biogenesis, has received considerable attention in recent years (25, 26). Here, immunofluorescence analysis showed that the fluorescence intensity of NRF1 in the nuclei of CD8^+^ T cells in the co-culture system treated with TD-139 was strongly increased compared with the non-inhibitor group (Figures 6B, C). Taken together, our findings suggest that galectin-3 mediates the down-regulation of mitochondrial complex III/IV genes by suppressing NRF1 nuclear translocation and transcriptional activation.

### ERK and/or Akt signaling pathway may be involved in CD8^+^ T cell mitochondrial dysfunction

To determine which signaling cascade galectin-3 functions through, phosphorylation levels of kinases involved in signaling pathways were examined using the proteome profiler array. After co-culture with or without inhibitor treatment, CD8^+^ T cells in the lower chamber were collected and proteins were extracted and then assayed. Quantification and statistical analysis of the results identified 25 differential proteins, among which the most significant down-regulated proteins in the inhibitor group were ERK, GSK-3, and EGFR. The markedly increased proteins included multiple transcription factors from STAT family, protein kinase B (Akt), and the molecular chaperone HSP60 (Figure 6D). It is interesting to note, the upregulation of Akt signaling and HSP60 is consistent with their known roles in promoting mitochondrial homeostasis. Furthermore, functional enrichment analysis showed that differential proteins were enriched in several pathways, including regulation of mitochondrial membrane potential and mitochondrion organization (Figure 6E). Our results showed that the downstream effect of galectin-3 might correlate with ERK activation and/or Akt suppression, which can be reversed by inhibitor treatment.

## Discussion

Understanding SARS-CoV-2-specific T cell responses is critical for viral clearance and development of effective strategies to prevent adverse clinical outcomes. The studies have shown that the decline and depletion of T cells are negatively correlated with the prognosis, especially in patients requiring intensive care (27, 28). Although T cell decrease has been described as a common symptom of severe patients, its mechanism was not fully understood. In this study, we utilized computational methods to integrate and analyze single-cell characterizations of BALF, combined with ex vivo experiments, and found that in severe COVID-19, galectin-3 secreted by lung epithelial cells acted on CD8^+^ T cells, downregulated mitochondrial complex gene transcription through NRF1, and caused cell cycle arrest and inhibition of CD8^+^ T expansion.

Naive CD8^+^ T cells can recognize SARS-CoV-2 antigens, and undergo clonal expansion and differentiation into effector CD8^+^ T cells (10). In our analysis at the single-cell level, CD8^+^ T cell cluster at the stage of severe/critical disease was found to have a proliferative phenotype with highly expressed MKI67 and PCNA; however, the proportion of CD8^+^ T cells in severe/critical patients decreased significantly compared with moderate patients, suggesting ineffective or failed clonal expansion. Meanwhile. the immune function of some cells in this group was partially impaired. Several previous studies have reported cells with similar phenotypes and called it a “proliferative-exhausted” phenotype (29, 30). Here, we further delved into the intercellular heterogeneity in this cluster, subdividing it into two subgroups and revealing the shared characteristics and differences between the two subpopulations in subsequent analyses.

TCR immune repertoire analysis showed that these two CD8^+^ T cell clusters were less clonal than corresponding clusters in moderate COVID-19 patients, indicating that these cells did not exert their proliferation function to increase CD8^+^ T cells in severe/critical patients as they should. Compared to moderate COVID-19 patients and healthy people, CD8^+^ T cells derived from severe/critical patients showed an increased proportion of cells in S phase and G2/M phase, indicating the blockade of cell cycle process. And the pseudotime analysis also suggested that these two populations were at another end of the developmental trajectory, which means these proliferative-exhausted cells may represent a divergent destination for naive CD8^+^ T cells. In particular, although sharing the same proliferative phenotype, cluster 10 showed obvious G2/M blockade with a high level of G2/M gene expression, while cluster 3 did not have a phase propensity. GSEA analysis revealed the activation of multiple negatively regulated cell cycle signaling pathways in cluster 10. These results suggested that these two CD8^+^ T cell clusters could be regulated diversely and lead to different transcriptional activities.

Energy metabolism plays a crucial part in the process of immune cell division and proliferation. During the activation upon antigen stimulation, the metabolic energy supply of T cells changes from a resting state to increased mitochondrial metabolism (31). Our single-cell analysis and experimental results confirmed that those cell-cycle-arrested CD8^+^ T lymphocytes had impaired mitochondrial biogenesis, in which the genes encoding subunits of electron transport chain (ETC) complexes were downregulated at the transcriptional level, leading to insufficient energy supply to meet the consumption required for cell division. This may partially explain why the specific T cell subsets in severe/critical patients expressed proliferation-related signatures but ultimately did not lead to T cell expansion — they never really get through the mitosis.

We herein reported that galectin-3 can regulate the cell cycle of CD8^+^ T cells by modulating their mitochondrial function. The study has shown that the SARS-CoV-2-infected cells trigger senescence-like cell-cycle arrest in the neighboring uninfected cells in a paracrine manner via virus-induced cytokine production (32). In the communication analysis of epithelial cells and CD8^+^ T cells, we noted the galectin signaling pathway. Galectin-3, a member of the β-galactoside-binding lectin family, was highly expressed by epithelial cells and had a stronger effect on cluster10. Articles have reported that the up-regulation of serum/plasma galectin-3 can be used as a prognostic biomarker in patients with COVID-19 (33, 34); however, its role in the disease process is not completely clear. In our study, we speculated that CD8^+^ T cells whose cell cycles were blocked in severe/critical COVID-19 patients may also be affected by infected epithelial cells and galectin-3 was the crucial ligand among them.

ORF3a is an accessory protein of SARS-CoV-2 and plays an important role in the immunopathogenesis of COVID-19. Several works have shown that ORF3a drives target cells into an inflammatory state by promoting apoptosis, inducing endoplasmic reticulum stress, and other mechanisms (22, 35). Lung epithelial cells were transfected with ORF3a to partially simulate the model of epithelial cell infection, and examine the host cells responding to virus infection. The results indicated that various pro-inflammatory factors and proteins including galectin-3, were upregrulated in the lung epithelial cells transfected with ORF3a. Our data further supported the profound effect of viral accessory protein on host cells.

While CD8^+^ T cells were co-cultured with lung epithelial cells transfected with ORF3a, the activation of nuclear respiratory factor 1 (NRF1) and the expression of mitochondrial complex genes were downregulated. The inhibitor of galectin-3 activity has been reported to ameliorate the mitochondrial damage in the heart of obese rats (36). As a nuclear respiratory factor, NRF1 can initiate the transcription of multiple mitochondrial respiratory chain-related genes. It was also discussed that NRF activators could be used to treat patient with COVID-19 (37, 38). In our co-culture system, NRF1 was significantly activated, and the expression of mitochondrial complex genes was recovered in the presence of TD-139.

The comparative analysis of galectin-3 inhibitor treated and nontreated groups showed multiple changes in different kinases, especially in ERK- and Akt-related signal pathways with opposing trends. The ERK 1/2 is the key to transmitting signals from surface receptors to the nucleus. It was found to inhibit mitochondrial biogenesis by phosphorylating different effector molecules in tumors and neurodegenerative diseases (39, 40). Consistent with the previous studies, galectin-3 inhibitor significantly reduced ERK phosphorylation in CD8^+^ T cells, accompanied by a rebound in mitochondrial complex gene transcription. In contrast, Akt phosphorylation was upregulated in our galectin inhibitor-treated group. Akt phosphorylation promoted the activation of PGC-1α, which is an important regulator of mitochondrial biogenesis to enhance respiratory capacity and maintain homeostasis (41). The regulation of these two kinases improved mitochondrial biogenesis and cell cycles. In addition, galectin-3 inhibitor up-regulated multiple transcription factors of the STATs family, which indicated in CD8+ T lymphocyte’s response to interferon signals and antiviral function was restored.

In conclusion, we identified a specific CD8^+^ T cell subset in severe/critical COVID-19 patients, which is characterized with abnormal proliferative gene expression and cell cycle arrest. Galectin-3 produced by infected epithelial cells participates in the regulation of CD8^+^ T cells by inhibiting the transcription of mitochondrial respiratory chain complex genes, resulting in mitochondrial biogenesis impairment and insufficient energy supply, eventually stalling the process of CD8^+^ T cell proliferation and expansion. Blocking galectin-3 may relieve the cell cycle arrest of CD8^+^ T cells in COVID-19 patients, and restore the proliferation ability and cell function of CD8^+^ T cells. As TD-139, a galectin-3 inhibitor, is already available as a treatment for idiopathic pulmonary fibrosis (IPF), our study provides new insight into the benefits of galectin-3 suppression in COVID-19 patients, which may save lives of severe/critically ill patients and also provide additional support to the lung function of patients who survive the chronic phase of COVID-19.

## Author Contributions

JZ and JH worked on the conception and design of the study. YW, CY and ZW analyzed the sequencing data including scRNA-seq and scTCR-seq, and visualized the figures. ZW, Yi Wang, QY, YL, YF, J Zhao and XZ performed the experiments and processed the experimental data. YW, Yi Wang, and ZW draft the initial manuscript. JZ and JH critically supervised the study and carefully revised the article. All authors listed contributed to the article and approved the final manuscript.

## Acknowledgments

We would like to thank Dr. Nevan Krogan’s lab and Addgene for their generosity in providing the plasmids used in this study. We gratefully acknowledge the financial support from the Department of Science and Technology of Hubei Province (Project# 2020BCB048 and 2022EHB035) and Hubei talent program (Project# 1180011).

